# Inter- and intra-specific variation in hair cortisol concentrations of Neotropical bats

**DOI:** 10.1101/2021.01.10.426004

**Authors:** Natalia I. Sandoval-Herrera, Gabriela F. Mastromonaco, Daniel J. Becker, Nancy B. Simmons, Kenneth C. Welch

**Affiliations:** Department of Ecology and Evolutionary Biology, University of Toronto, Ontario, Canada; Department of Biological Sciences, University of Toronto Scarborough, Ontario, Canada; Reproductive Sciences, Toronto Zoo, Ontario, Canada; Department of Biology, University of Oklahoma, Norman, OK 73019; Department of Mammalogy, Division of Vertebrate Zoology, American Museum of Natural History, New York, NY

## Abstract

Quantifying hair cortisol has become popular in wildlife ecology for its practical advantages for evaluating health. Before hair cortisol levels can be reliably interpreted however, it is key to first understand the intrinsic factors explaining intra- and interspecific variation. Bats are an ecologically diverse group of mammals that allow studying such variation. Given that many bat species are threatened or have declining populations in parts of their range, non-invasive tools for monitoring colony health and identifying cryptic stressors are needed to efficiently direct conservation efforts. Here we describe intra- and interspecific sources of variation in hair cortisol levels in 18 Neotropical bat species from Mexico and Belize. We found that fecundity is an important ecological trait explaining interspecific variation in bat hair cortisol. Other ecological variables such as colony size, roost durability, and basal metabolic rate did not explain hair cortisol variation among species. At the individual level, females exhibited higher hair cortisol levels than males, and the effect of body mass varied among species. Overall, our findings help validate and accurately apply hair cortisol as a monitoring tool in free-ranging bats.

## Introduction

Free-living animals face multiple natural and anthropogenic challenges that threaten their survival and thus are of considerable interest to ecophysiologists concerned with the study of effects of stress on vertebrates. One of the most extensively studied processes associated with response to stressors (biotic or abiotic environmental factors that disrupt homeostasis; Schulte, 2014) is the release of glucocorticoid (GC) hormones (Creagh and Brendan Delehanty, 2013; MacDougall-Shackleton *et al.*, 2019). GCs are known to facilitate the mobilization of energy required to cope with stressors and, during normal conditions, play a key role in regulating growth, circadian activity, and energy metabolism (review in Landys *et al.*, 2006). Levels of GCs are commonly employed as a biomarker of health or relative condition (Sapolsky *et al.*, 2000; Wikelski and Cooke, 2006; Pearson Murphy, 2007; Busch and Hayward, 2009). GC secretion is a well-conserved process across vertebrates and involves activation of the hypothalamic-pituitary-adrenal (HPA) axis and release of GCs from the adrenal glands to the blood stream (Norris and Carr, 2013). In mammals, the primary GC is cortisol, which induces a cascade of events to maintain homeostasis at multiple target tissues (Pearson Murphy, 2007; Boonstra, 2013). An acute increase in GC levels can benefit an individual’s survival (e.g., by allocating energy in defense and escape) yet if adverse conditions remain, continuously elevated GCs in circulation can become pathological, causing immune suppression, neuronal cell death, and reproductive impairment(Sapolsky *et al.*, 2000; Tilbrook, 2000; Wingfield and Romero, 2011; Hing *et al.*, 2016).

Although many of the environmental challenges that wild populations experience are chronic (e.g., prolonged food deprivation, climate change, habitat disturbance, pollution), studies of stress physiology have focused on detecting acute stress by looking at GC levels in blood, urine, and feces (Sheriff *et al.*, 2011; Creagh and Brendan Delehanty, 2013). The rapid turnover of these tissues, however, only gives short-term information of HPA activity over periods of hours or days (Sheriff *et al.*, 2011) which may not be an appropriate time scale. Assessment of cortisol in tissues with slower turnover rates, such as hair, may reflect circulating cortisol levels over longer periods of several weeks or even months, which is the time scale over which chronic environmentally-induced stress would be expected to occur (Davenport *et al.*, 2006; Macbeth *et al.*, 2010; Ashley *et al.*, 2011; Mastromonaco *et al.*, 2014). Cortisol is incorporated into developing hairs from the blood stream during periods of active hair growth, allowing researchers to retrospectively examine cortisol production at the time that a stressor or stressors were faced (Davenport *et al.*, 2006; Pragst and Balikova, 2006). Hair can be collected in a relatively non-invasive manner, is usually easily accessible in relatively large amounts, and is easy to store and transport, all of which make it particularly useful for wildlife studies, especially those involving threatened or endangered species (Koren et al., 2002; Macbeth et al., 2010; Macbeth et al., 2012). Hair cortisol levels are not likely affected by stress induced by capture and/or handling, which is one of the main limitations of blood GC analysis (Russell *et al.*, 2012). A single sample of hair can also provide complementary and valuable information about ecology and behavior, including diet and movement (e.g., using stable isotope analyses; Fraser *et al.*, 2010; Sullivan *et al.*, 2012; Voigt *et al.*, 2012; Oelbaum *et al.*, 2019), condition (e.g. nutrition; Montillo *et al.*, 2019), toxicant exposure (Hernout *et al.*, 2016; Becker *et al.*, 2018), and molecular identification (Magioli *et al.*, 2019), opening possibilities for more integrative studies. However, analyses of hair samples can be challenging. Despite being a very promising tool for assessing wildlife health, quantifying hair cortisol is a method that has limitations, though these are largely based on lack of detailed knowledge of patterns of hair grown (Meyer and Novak, 2012; Russell *et al.*, 2012; Sharpley *et al.*, 2012). For example, the exact time scale reflected in any given sample will depend on the rate of hair growth and moulting patterns; this information is unknown for most species, which makes the time window being evaluated unclear (Koren *et al.*, 2002; Fourie *et al.*, 2016). Moreover, rates of cortisol incorporation to the hair shaft are known to differ across body regions and among species (Sharpley *et al.*, 2012; Acker *et al.*, 2018; Lavergne *et al.*, 2020). Nevertheless, hair cortisol levels offer a potentially powerful tool for assessing relatively long-term stress levels in mammals.

Hair cortisol and its correlation with natural and anthropogenic stressors has been explored for different wild mammals, including rhesus monkeys (*Macaca mulatta*; Dettmer et al., 2012), grizzly bears (*Ursus arctos*; Macbeth et al., 2010), reindeer/caribou (*Rangifer tarandus*; Ashley et al., 2011), lynx (*Lynx canadensis*; Terwissen *et al.*, 2013), mongoose (*Herpestes ichneumon*; Azevedo *et al.*, 2019), and snowshoe hares (*Lepus americanus*; Lavergne *et al.*, 2020). Although most of these studies support hair cortisol as an informative measure of central HPA activity, they also identified intrinsic factors such as age, sex, reproductive stage, and social status that modulate GCs levels in different contexts (Wingfield and Romero, 2011; Crespi *et al.*, 2013; Hau *et al.*, 2016). Not accounting for these intrinsic sources of variation in GC levels may lead to incorrect or misleading estimates of the effects of stressors on individual fitness and population health (Sapolsky *et al.*, 2000; Reeder and Kramer, 2005; Busch and Hayward, 2009; Wingfield and Romero, 2011; Kalliokoski *et al.*, 2019).

Ecological traits such as diet, fecundity, and lifespan, as well as phylogenetic relatedness, have been proposed to explain differences in baseline cortisol levels in wild species (Wingfield and Romero, 2011; Patterson *et al.*, 2014). Evolution of different life-history strategies are also thought to have led to different adaptations in HPA activity modulation so as to maximize individual fitness within species (Bonier *et al.*, 2009; Bonier and Martin, 2016). Bats are a very ecologically diverse group comprising over 1,400 species that live in most terrestrial ecosystems and have a wide variety of diets, use many different roost types, and have many different social systems (Kunz and Fenton, 2005; Dumont *et al.*, 2012; Gunnell and Simmons, 2012; Simmons and Cirranello, 2020). This diversity provides the opportunity to study the ecological correlates of cortisol levels among phylogenetically related species with different life-history traits. Few ecological correlates of GCs have been evaluated simultaneously in mammalian groups in the context of cortisol studies, and fewer studies have further related cortisol levels to life-history traits across multiple species from a single mammalian clade. Among bats, variation in hair cortisol levels associated with seasonal food availability has been studied in two species with contrasting diets, *Carollia perspicillata* and *Desmodus rotundus* (Lewanzik *et al.*, 2012), but no other comparative studies have been conducted within this order. Furthermore, little is known about the modulation of the stress response in bats, despite Chiroptera being the second most speciose order of mammals.

Bat populations are declining worldwide due to ongoing habitat destruction and land use changes, increased interaction with human environments and associated threats including wind turbine fatalities, hunting and targeting killing, pesticide exposure, and emerging infectious diseases such as white-nose syndrome (O’Donnell, 2000; Mickleburgh *et al.*, 2002; Kunz *et al.*, 2007; Frick *et al.*, 2010; Racey, 2013; Voigt and Kingston, 2015). Because many bat species are threatened or have declining populations in parts of their range (IUCN Red List of Threatened Species, 2020), non-invasive tools to monitor colony health and identify cryptic stressors are critically needed to efficiently direct conservation efforts. It is essential to investigate the factors influencing baseline GCs to properly detect elevated cortisol levels due to long-term stressors.

In this study, we describe intra- and interspecific sources of variation in baseline hair cortisol levels in bats, which contributes to better understanding the potential for hair cortisol to be an indicator of HPA activity in this taxon. We hypothesize that interspecific variation in hair cortisol of bats will be greater than intraspecific variation, and that such heterogeneity will be best explained by ecological traits directly related to energy expenditure, such as basal metabolic rate (BMR), dietary guild, foraging behavior, and roost durability. We expect that species with high energetic demands or less predictable energy acquisition (e.g., less reliable food sources) will have higher hair cortisol. Specifically, we predict that: 1) a positive relationships between BMR and hair cortisol; 2) bats that feed on fruit and nectar - which are energy-rich and readily available - will have lower hair cortisol; 3) bats that actively hunt prey during flight, such aerial hawkers, will have higher GC levels owing to greater energetic demands compared to gleaners that can hunt from perches (Norberg and Rayner, 1987; Fenton, 1990); and 4) species using more ephemeral day roosts (e.g. foliage or crevices under exfoliating bark), will have higher hair cortisol than species using more stable structures (Kunz and Fenton, 2005).

## Material and methods

### Study Sites

We sampled bats from northern Belize (Orange Walk District) and two locations in Mexico (Colima and Chihuahua States). In each region, we sampled sites with different levels of habitat fragmentation and agricultural intensity. We used the global Human Modification Index (HMI; Kennedy *et al.*, 2019) as a standardized measure of disturbance, using a 5 km buffer around each collection site. The HMI is a cumulative measurement with possible values between 0 (no disturbance) and 1 (highest disturbance) that includes transportation, human settlement, agriculture, extractive activities, and electric infrastructure (Kennedy *et al.*, 2019). Sites were classified as low (0 median HMI ≤ 0.10), moderate (0.10 < median HMI ≤ 0.40), high (0.40 < median HMI ≤ 0.70), and very high (0.70 < median HMI ≤ 1.00). At all sites, bats were captured from 18:00 to 22:00 hrs using mist nets and from 18:00 to 5:00 using harp traps (only in Belize) set along flight paths. Bats sampled during the day were captured in their roosts, mainly caves, using hand nets. We recorded sex, size (body mass [g],forearm length [mm]), and reproductive stage (pregnant, active, inactive; Kunz and Parsons, 2009).

In Colima (west central Mexico) in March 2019 (dry season), we sampled bats roosting in three caves surrounded by different levels of disturbance: Don Pancho Cave (moderate disturbance, HMI=0.38), El Salitre Cave (high disturbance, HMI=0.44) and Coquimatlán Cave (high disturbance, HMI=0.57; Fig 1). Don Pancho Cave, is located on San Agustin island, 1 km away from the coast of Chamela Bay, Jalisco (19.5353°N, −105.0881°W). El Salitre Cave is near Los Ortices village, Colima (19.083330°N,-103.726667°E). La Fábrica Cave is 6.4 km SW of Coquimatlan town, Colima (19.1513°N,-103.8353°W). We refer here to these locations collectively as central Mexico. we also sampled bats foraging close to pecan nut croplands near the town of Jimenez, Chihuahua (northern Mexico). This region is entirely dedicated to the production of pecan nuts with thousands of squared kilometers of cultivated land (Orona Castillo *et al.*, 2018). We visited one that farms using organic practices and another that farms with intensive use of pesticides. However, the estimated HMI index was the same for the two sites (HMI=0.49, high disturbance). We collected hair samples from three bat species (*Antrozous pallidus, Tadarida brasilensis*, and *Myotis velifer*) at both northern Mexico sites.

**Figure 1.**
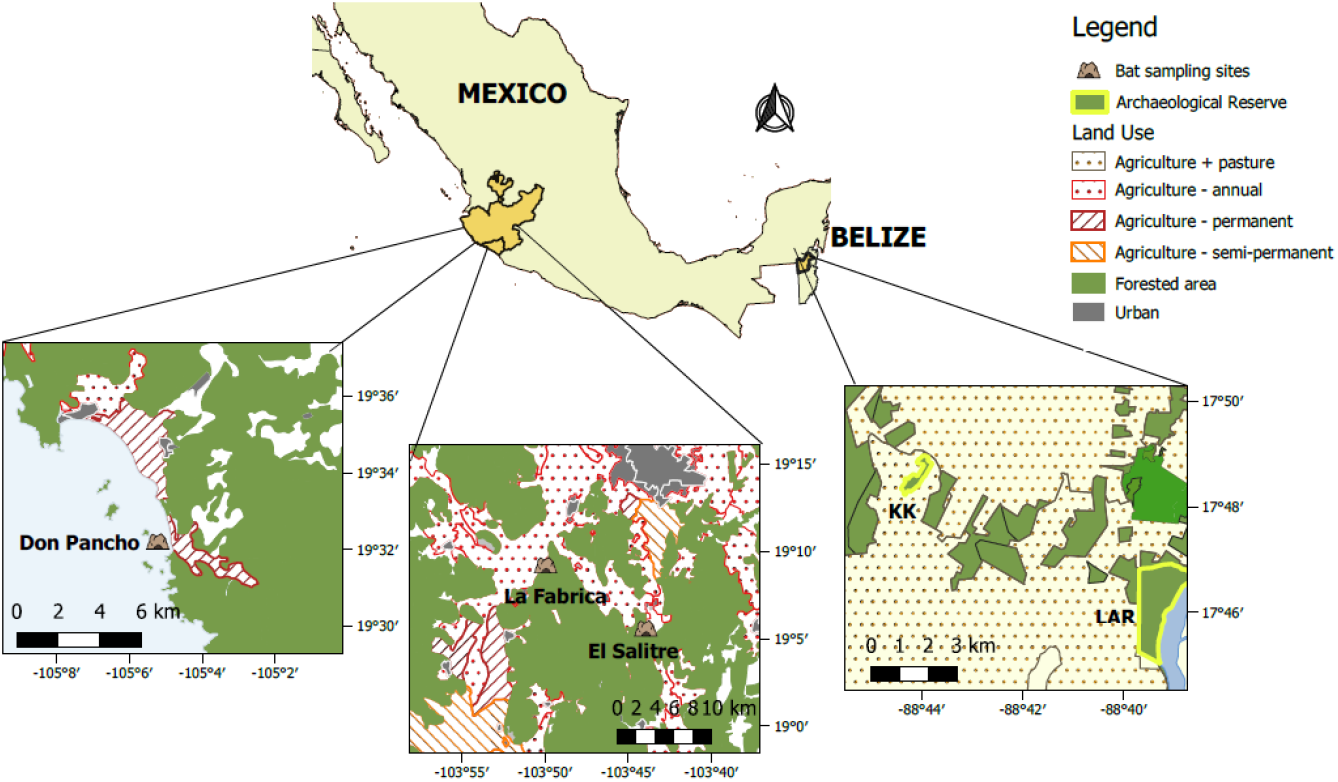
Sampling sites in central Mexico and Belize, showing the use of land in the surrounding areas. Sources: Sistema Nacional de Información Estadística y Geográfica de Mexico (INEGI, 2013) and Biodiversity and Environmental Resource Data System for Belize (BERDS).

**Figure 2.**
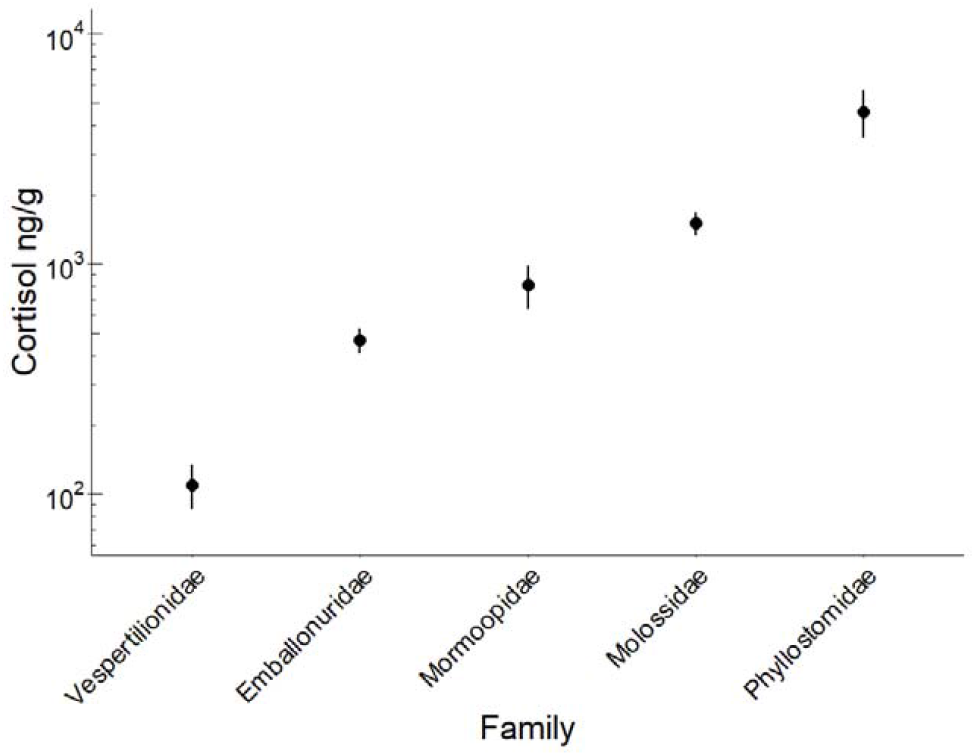
Cortisol concentration in hair samples from 18 Neotropical bat species grouped by family. Error bars represent the Standard Error of the mean. The y axis is in log scale.

Our field sites in Belize consisted of two forest patches of very different size located approximately 10 km apart and separated by a heterogeneous, largely agricultural landscape. Lamanai Archaeological Reserve (LAR) is a protected secondary semi-deciduous forest of 450 ha with a high canopy and with relatively low disturbance (HMI=0.17) (Herrera *et al.*, 2018). In contrast, the Ka’Kabish archeological site (KK) is a small remnant forest patch of about 45 ha surrounded by cattle pastures and local croplands (Fig 1). Although the landscape in Belize is apparently disturbed and highly fragmented, agricultural activity and urban development is not as intense as the field sites in Mexico, which is reflected in their moderate HMI scores (LAR: 0.17; KK: 0.18). We collected hair samples from 13 different species (Table 1) in April 2018 and 2019 (dry season) at Belize sites.

**Table 1.** Species-level ecological traits and hair cortisol data for 18 Neotropical bat species.

| Family | Species | Region | N | Mass(g) | Dietary niche | Foraging style | WAR | Roost durability | Fecundity (litter/yr) | Colony size | Lifespan (years) | Cortisol (ng/g) |  |  |
| --- | --- | --- | --- | --- | --- | --- | --- | --- | --- | --- | --- | --- | --- | --- |
| | | | | Mean ± SD | | | | | | | | Mean | ± | SD |
| Emballonuridae | *Rhynchonycteris naso* | BZ | 14 | 4.54 ± 1.47 | Animalivory | Aerial forager | 6.5 | 2 | >1 | Small | 5 | 639,57 | ± | 392,10 |
| | *Saccopterix billineata* | BZ | 20 | 6.79 ± 1.12 | Animalivory | Aerial forager | 6.1 | 2,4 | 1 | Small | 5 | 347,28 | ± | 232,01 |
| Molossidae | *Molossus nigricans* | BZ | 5 | 35.00 ± 2.35 | Animalivory | Aerial forager | 11.1 | 4 | 1 | Large | 10 | 227,80 | ± | 123,54 |
| | *Tadarida brasiliensis* | NMX | 23 | 10.73 ± 0.63 | Animalivory | Aerial forager | 8.6 | 5,3 | 1 | Large | 10 | 1782,87 | ± | 751,07 |
| Mormoopidae | *Pteronotus mesoamericanus* | BZ | 26 | 17.08 ± 2.06 | Animalivory | Aerial forager | 6.7 | 5,6 | 1 | Large | 10 | 218,79 | ± | 76,48 |
| | *Pteronotus mexicanus* | CMX | 35 | 13.37 ± 1.18 | Animalivory | Aerial forager | 6.7 | 5,6 | 1 | Large | 10 | 1246,96 | ± | 1663,56 |
| Phyllostomidae | *Desmodus rotundus* | BZ | 22 | 28.01 ± 3.54 | Animalivory | Gleaner | 6.7 | 4,5 | >1 | Large | 15 | 395,00 | ± | 226,92 |
| | *Glossophaga soricina* | BZ | 19 | 10.33 ± 1.87 | Phytophagy | Gleaner | 6.4 | 4,6 | >1 | Medium | 10 | 99,41 | ± | 117,86 |
| | *Leptonycteris yerbabuenae* | CMX | 18 | 22.26 ± 1.72 | Phytophagy | Gleaner | 7 | 5,6 | 1 | Large | 15 | 24614,50 | ± | 14780,52 |
| | *Lophostoma evotis* | BZ | 1 | 19.00± | Animalivory | Gleaner | 5.3 | 3 | 1 | Small | 20 | 3046,00 | ± | |
| | *Macrotus waterhousii* | CMX | 23 | 14.31 ± 2.63 | Animalivory | Gleaner | 5.8 | 5,6 | 1 | Medium | 10 | 616,37 | ± | 415,85 |
| | *Mimon cozumelae* | BZ | 3 | 25.00 ± 0 | Animalivory | Gleaner | 8.3 | 2,8 | 1 | Small | 20 | 2000,67 | ± | 1284,91 |
| | *Sturnira parvidens* | BZ | 17 | 14.61 ± 1.55 | Phytophagy | Gleaner | 6.5 | 4 | >1 | Small | 20 | 635,97 | ± | 379,79 |
| | *Trachops cirrhosus* | BZ | 3 | 31.50 ± 0.50 | Animalivory | Gleaner | 6.3 | 3,9 | 1 | Medium | 10 | 147,10 | ± | 22,36 |
| Vespertilionidae | *Antrozous pallidus* | NMX | 10 | 15.80 ± 1.70 | Animalivory | Gleaner | 6.5 | 4 | >1 | Small | 10 | 42,79 | ± | 17,46 |
| | *Eptesicus furinalis* | BZ | 7 | 7.57 ± 0.45 | Animalivory | Aerial forager | 6.2 | 3 | >1 | Large | 20 | 36,66 | ± | 40,54 |
| | *Lasiurus ega* | BZ | 2 | 12.00 ± 4.24 | Animalivory | Aerial forager | 7.9 | 2,5 | >1 | Small | 15 | 142,83 | ± | 18,14 |
| | *Myotis velifer* | NMX | 11 | 8.98 ± 1.02 | Animalivory | Aerial forager | 6.7 | 2,5 | 1 | Large | 10 | 211,65 | ± | 170,79 |

\*Site: capture site BZ: Belize, NMX: Northern Mexico, CMX: Central Mexico; N: Number of hair sample analyzed for cortisol; Species were assigned to two dietary niches: Animalivory (insectivorous, sanguinivorous and carnivorous) and phytophagy (frugivorous and nectarivorous species). Roost category assignment followed Patterson et al., 2011. WAR: Wing Aspect Ratio .

### Ethical statement

Field procedures followed guidelines for safe and humane handling of bats published by of the American Society of Mammalogists (Sikes and Bryan, 2016) and were approved by the Institutional Animal Care and Use Committees of the University of Georgia (A2014 04-016-Y3-A5), University of Toronto (20012113), and American Museum of Natural History (AMNHIACUC-20180123). Fieldwork was authorized by the Belize Forest Department under permits WL/2/1/18(16) and WL/1/19(06). Sample collection in Mexico was approved under the permit #FAUT-0069.

### Sample collection

We trimmed a single hair sample (3-10 mg) from the scapular region on the back of each bat and stored resulting samples individually in 1-2 ml sample tubes. The amount of hair removed from each bat depended on the hair density of each species. The hair shaft was carefully cut close to the root avoiding removing skin or follicle tissue. From pilot analyses, we determined a minimum amount of 3 mg of hair was necessary to obtain values around 50% binding on the standard curve thereby accurately estimating cortisol concentration in the sample.

### Extraction and quantification of cortisol

Hair samples were processed and analyzed at the Endocrinology Laboratory at the Toronto Zoo following methods described by Acker *et al.*, 2018. Each hair sample was spread apart and weighed in a 7 mL glass scintillation vial. To avoid contamination with other biological fluids =, all hair samples were washed with 100% methanol by vortexing in a tube for 10 s and immediately removing the methanol using a pipettor. Immediately thereafter, 80% methanol in water (v:v) was added to each sample, at a ratio of 0.005 g/mL. Samples were then mixed for 24 h on a plate shaker (MBI Orbital Shaker; Montreal Biotechnologies Inc., Montreal, QC, Canada). After 24 hrs the vials were centrifuged for 10 min at 2400g. The supernatants were pipetted off into clean glass vials and dried down under air in a fume hood. The dried extracts were stored at −20°C until analysis.

Samples were brought to room temperature prior to analysis. Reconstitution of the desiccated extracts was done by adding phosphate buffer and vortexing for 10 s. Belize samples were reconstituted neat (i.e. evaporated 150ul and reconstituted with 150ul) and Mexico samples were reconstituted as follows: four species were neat, two species diluted 1:5 and one species diluted 1:50 in phosphate buffer (Andreasson *et al.*, 2015). Cortisol concentrations were determined using an EIA (R4972, C. Munro, University of California, Davis); antibody and HRP dilutions were 1:10,200 and 1:33,400, respectively. All samples were centrifuged for 1 min at 1200g immediately prior to dispensing onto the microtiter plate. Results are presented as nanograms of cortisol per gram of hair.

### Species ecological traits

We compiled data on ecological traits considered relevant to cortisol mobilization from previously published literature and databases. Values for traits are species-level averages and may not reflect specific values at these sites (Table1). Data on Basal Metabolic Rate (BMR) was extracted from the literature (Cruz-Neto *et al.*, 2001; Genoud *et al.*, 2018) and when not available (n=2) the following formula was used for the estimation: *ln BMR* = 0.744 × In *mass*(*in g*) + 1.0895 (Speakman and Thomas, 2003).Information on diet, foraging style, percentage of invertebrates in the diet, and fecundity was extracted from the Elton Traits, PanTHERIA, and Amniote Life History databases (Myhrvold et al., 2015; Wilman et 206 al., 2014). We collapsed variation in diet into two dietary niches: phytophagy (including nectarivores and frugivores) and animalivory (insectivores and carnivores) because many bat species in our study have diets that combine more than one food source within these categories (Fenton *et al.*, 2001; Kunz and Fenton, 2005; Reid, 2009; Oelbaum *et al.*, 2019). We also considered the percentage of invertebrates in the diet of the animalivorous bats, which can vary significantly among species. Because foraging behavior is a complex and plastic trait, we simplified this variable into two categories: aerial foragers (i.e., hawkers) and gleaners (including species that glean plant products like fruit as well as insects) since these behaviors may reflect differences in energetic demands associated with foraging (Herrera *et al.*, 2018). Because wing morphology can strongly influence the energetic costs of flight, we also included the mean wing aspect ratio for each species (Norberg and Rayner, 1987; Bullen *et al.*, 2014). Fecundity was defined as the annual average fecundity (litter size × number of litters per year). We estimated roost durability following the methods of Patterson et al. (2007), where 1 indicates the most ephemeral and least protected roost types (e.g. rolled leaves and foliage) and 6 indicates the most permanent and protected roost types (e.g. caves). For species known to multiple use different kinds of roost, intermediate ranks were calculated, weighing roost categories according to the relative frequency of use reported in the literature (Schneeberger *et al.*, 2013). Lifespan was drawn from the Animal Ageing and Longevity database(AnAge: The Animal Ageing and Longevity Database, 2020) and DATLife (DATLife Database. Max-Planck Institute for Demographic Research (Germany), 2020). For many of the species in these databases, longevity estimates are based on captive animals, which likely overestimates life expectancy in the wild. Because bats of a single species may live in colonies of varying sizes, and most values on colony size are reported in ranges in the literature, we classified maximum colony sizes reported for each species as small (1-50) medium (50-500) or large (>500) *sensu* Santana *et al.* (2011).

### Data analysis

We first used phylogenetic generalized least squares (PGLS) models to evaluate the effect of species-level ecological variables on hair cortisol concentrations while accounting for bat phylogenetic relatedness. We used the *rotl* and *ape* packages in R to extract the bat phylogeny from the Open Tree of Life and calculate branch lengths with Grafen’s method (Paradis *et al.*, 2004; Michonneau *et al.*, 2016). We first fit a null PGLS model (intercept only) using the *nlme* package to estimate phylogenetic signal as Pagel’s *λ* (Pagel, 1999). We next fit a PGLS model with bat family as the predictor to assess broad taxonomic patterns in hair cortisol. We then fit 15 PGLS univariate models with, dietary niche, foraging behavior, roost durability, fecundity, lifespan, and colony size as predictors. We also fit fivemultivariate PGLS models including: BMR + body mass, niche + fecundity, niche + % invertebrates, niche + lifespan + fecundity, and niche + fecundity + colony. We compared PGLS models with Akaike information criterion corrected for small sample sizes (AICc) and assessed fit with an adjusted R^2^ (Burnham and Anderson, 2002). All PGLS models included weighting by sampling variance to account for variable sample sizes per species (Pennell, 2015).

We used generalized linear models (GLMs) to determine which individual- and habitat-level factors influence hair cortisol for each bat species. We first evaluated the relationship between body mass and hair cortisol separately for each species. Next, we ran species-specific GLMs including sex, reproductive stage, and site disturbance and predictors. Not all covariates were tested for all species due to sample size restrictions. Total sample size and balanced sample sizes among levels were considered to select the number of covariates to include in the model for each species. We included disturbance in GLMs only for species present in more than one site (*P. mesoamericanus, P. mexicanus, M. waterhousii, T. brasiliensis, G. soricina, D. rotundus*) since disturbance was treated as constant within sites. The only genus sampled in both Belize and Mexico was *Pteronotus*. The two species *P. mesoamericanus* (Belize) and *P. mexicanus* (Mexico) represent lineages considered conspecific until a few years ago, but are now thought to represent distinct species that diverged very recently based on molecular and morphometric evidence (Pavan and Marroig, 2016). Because their phenotypes and ecology are still very similar, we treated these as conspecific to test if there were differences in hair cortisol between representatives from the two regions (Mexico and Belize). Tukey post-hoc tests were conducted for significant covariates. We compared effect sizes across bat species by evaluating the degree of overlap in 95% confidence interval for each GLM coefficient. All analyses used the natural logarithm of hair cortisol as the response variable and assumed Gaussian errors. We report data as mean ± SD, unless otherwise noted.

## Results

### Ecological and evolutionary predictors of hair cortisol

We analyzed 262 hair samples from 18 different bat species representing five families in Belize and Mexico (Table 1). Hair cortisol concentration across species varied by four orders of magnitude, ranging from 36.6 ± 40.5 ng/g in *Eptesicus furinalis* to 24614 ± 14780 ng/g in *Leptonycteris yerbabuenae* (Table1). Mean hair cortisol did not differ much across families (F = 1.84, p = 0.02, R^2^ = 0.16), and family was not a good predictor of cortisol levels when compared to ecological and life history traits (Table 2). We did not find strong phylogenetic signal in species-level mean hair cortisol (Pagel λ=0). Using ecological traits, mean hair cortisol was best predicted by a model including both dietary niche and fecundity, although only fecundity had a significant effect (F_2,15_ =5.51; p=0.01; R^2^=0.34; Table 2). Annual fecundity explained 24% of the variance in Neotropical bat mean hair cortisol. Species reported to have more than one pup per year had significantly lower cortisol than bats having only one pup per year (F_1,16_ =6.22; p=0.02; Fig 3). Other ecological traits including roost durability, foraging strata, and colony size were uninformative predictors (Table 2).

**Figure 3.**
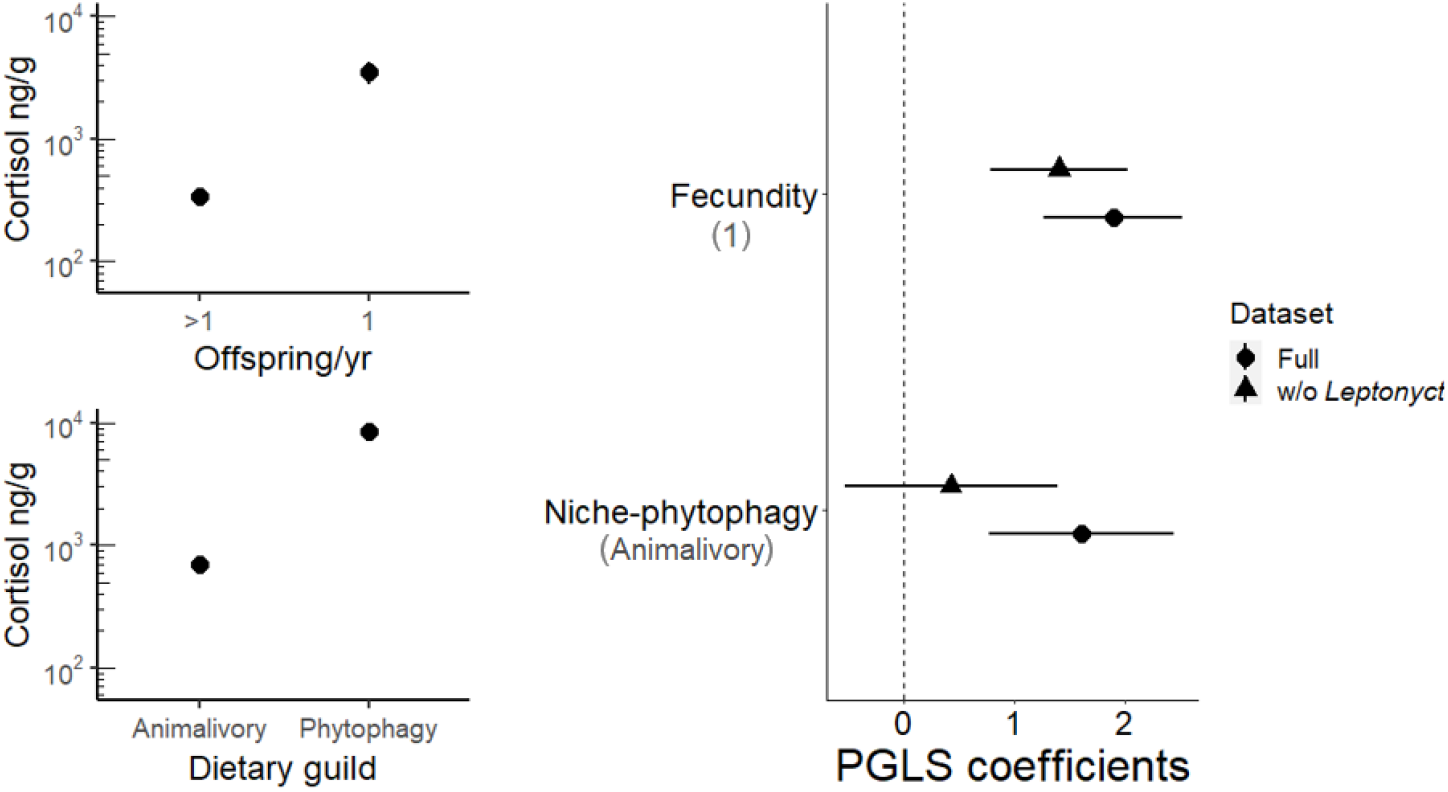
*Left:* Cortisol concentration in hair samples from 18 Neotropical bat species according to diet and annual fecundity. Y axes are in log scale.*Top*: Mean hair cortisol by number of offspring per year; *Bottom:* Hair cortisol by dietary niche (animalivory or phytophagy). *Right:* Differences in parameter estimates for the PGLS model with and without *Leptonycteris yerbabuenae.* Bars indicate the 95% confidence interval. The reference values for each variable of the model are listed in parentheses.

**Figure 4.**
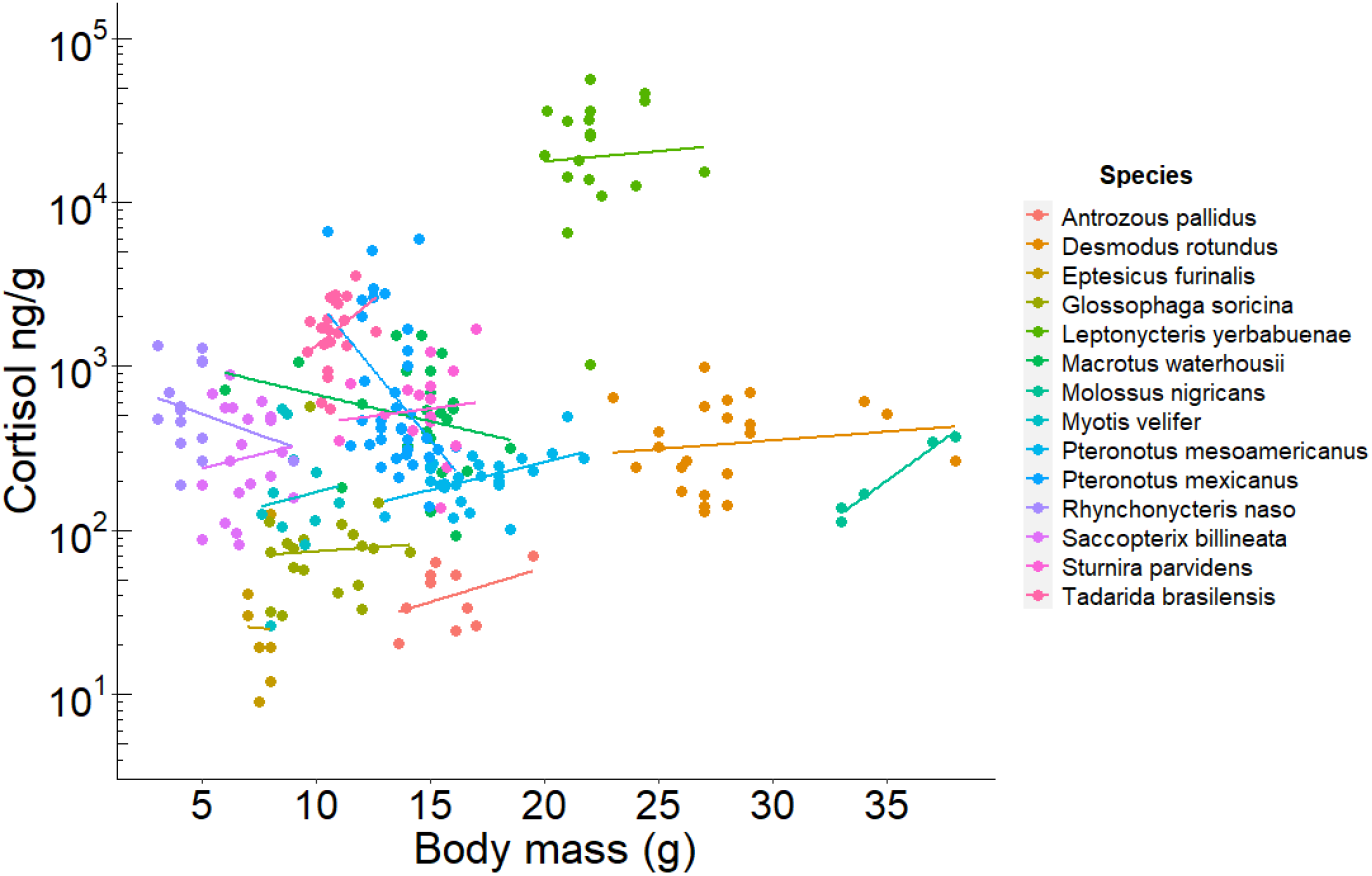
Relationship between hair cortisol concentration and body mass for each Neotropical bat species. Lines represent the GLM fit for each species. The y axis is in log scale.

**Table 2.** PGLS models predicting hair cortisol (ln transformed) in Neotropical bats. Models are ranked by ΔAICc with the number of coefficients (k), Akaike weights (wi), and the adjusted R^2^.

| Model structure | df | $\Delta\text{AICc}$ | $w_i$ | $R^2$ |
| --- | --- | --- | --- | --- |
| ~ niche+ fecundity | 2 | 0 | 0.412 | 0.34 |
| ~ fecundity | 4 | 1.08 | 0.247 | 0.24 |
| ~ niche+ fecundity+ foraging style | 5 | 3.30 | 0.081 | 0.30 |
| ~ 1 (intercept only) | 6 | 4.12 | 0.054 | 0 |
| ~ niche+ fecundity+ lifespan | 1 | 4.38 | 0.048 | 0.46 |
| ~ roost durability | 2 | 5.64 | 0.025 | 0 |
| ~niche | 3 | 5.68 | 0.024 | 0.01 |
| ~BMR+ body mass | 3 | 6.00 | 0.021 | 0 |
| ~sample | 2 | 6.39 | 0.017 | 0.33 |
| ~foraging style | 2 | 6.67 | 0.015 | 0 |
| ~colony size | 4 | 8.01 | 0.007 | 0.37 |
| ~niche + invertebrate% | 3 | 8.35 | 0.006 | -0.03 |
| ~BMR + foraging style | 3 | 8.92 | 0.005 | -0.08 |
| ~family | 5 | 9.12 | 0.004 | 0.16 |
| ~foraging style + WAR | 3 | 9.15 | 0.004 | -0.1 |
| ~lifespan | 4 | 10.54 | 0.002 | -0.06 |

The lesser long-nosed bat (*Leptonycteris yerbabuenae*) showed particularly high hair cortisol (26,6 ± 14, 8 ng/g). Because the high values of this species could bias inter-species comparisons, we assessed the sensitivity of our top models by excluding *L. yerbabuenae*. In Figure 3, we show how the coefficients from the PGLS top models with and without this species. In both cases, fecundity was the best ecological predictor of hair cortisol regardless of including *L. yerbabuenae* in the analyses.

### Individual-level analyses of bat hair cortisol

When investigating intraspecific variation, we found positive relationships between body mass and hair cortisol in two species: *Pteronotus mesoamericanus* (F_1,24_ = 7.34; p = 0.010; R^2^ = 0.23) and *Molossus nigricans* (F_1,4_ = 96.52; p = 0.002; R^2^ = 0.96; Fig 5). The opposite trend was found in *Pteronotus mexicanus* where heavier bats presented lower cortisol (F_1,33_ = 7. 97; p<0.01; R^2^ = 0.19). For *Desmodus rotundus*, only sex was a significant predictor of hair cortisol (F_1,19_ = 4.39; p = 0.04; R^2^ = 0.19). Male vampire bats had significantly lower hair cortisol than females (t_20_ = 2.09; p = 0.02). Similarly, variation in hair cortisol in the mustached bat (*P. mesoamericanus*) and the mastiff bat (*Molossus nigricans*) was explained only by sex, with males having lower concentrations than females (t_4_ = 2.68; p = 0.01 and t_24_ = −6.373; p = 0.01, respectively; Fig.6). When treating *Pteronotus mesoamericanus* (Belize) and *P. mexicanus* (Mexico) as one species, we found differences in hair cortisol between the two populations. Bats from Mexico had higher cortisol than their counterparts in Belize (F_1,59_ = 29.88; p<0.01; R^2^ = 0.33). Within Mexico, hair cortisol in *P. mexicanus* was explained by site disturbance (F_2,31_ = 72.35; p<0.001): bats roosting in Don Pancho cave (San Agustin island), a site with moderate disturbance (HMI = 0.38), showed significantly higher hair cortisol than bats roosting in El Salitre and La Fabrica caves in Colima (t_20_=9.94 p<0.01; t_21_=-10.29,Fig 1). There was no effect of sex (t_33_ = −1.15p>0.31) or females’ reproductive stage (F_4,14_ 3.26; p = 0.06) on hair cortisol in *P. mexicanus*. For other species such as *Eptesicus furinalis* (F_1,12_ = 2.451; p = 0.64), *Leptonycteris yerbabuenae* (F_1,17_=0.52; p=0.94), *Saccopteryx billineata* (F_2,17_ =0.18; p = 0.83), *Rhynchonycteris naso* (F_2,11_ = 2.60; p = 0.12), *Glossophaga soricina* (F_3,15_ = 0.13; p = 0.94), *Macrotus waterhousii* (F_2,18_ = 0.24; p = 0.78), *Sturnira parvidens* (F_2,14_ = 0.2052, p = 0.81), and *Antrozous pallidus* (F_2,9_ = 0.506;p = 0.68), none of the traits examined were informative predictors of hair cortisol levels.

**Figure 5.**
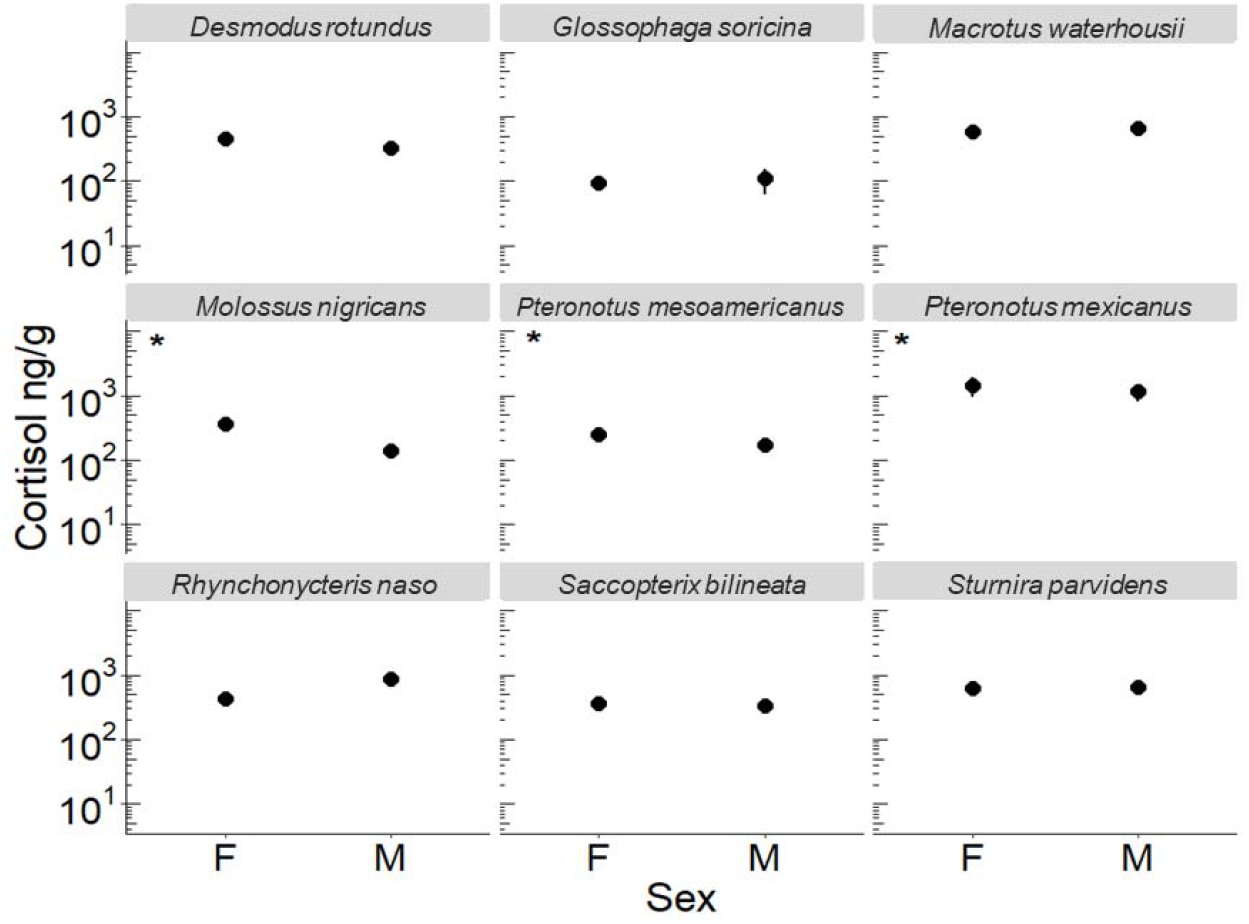
Mean hair cortisol concentration by sex for nine species of Neotropical bat species. Asterisks indicate species for which sex has a significant effect. Effects only shown for species with balanced sample sizes per sex. Error bars represent the standard error of the mean. Y axes are in log scale.

## Discussion

Analysis of hair cortisol has become a popular method to study long-term stress in wild animals, offering several practical advantages (e.g minimally invasive collection, easy sample storage and transport). An accurate interpretation of cortisol levels attributed to stress, however, requires a good understanding of the intrinsic and extrinsic drivers of baseline variation. Factors influencing hair cortisol in bats must be identified before hair cortisol can be used as a conservation tool to assess effects of environmental conditions on bat population health. In this study, we present the first quantification of hair cortisol in bats and describe relationships between hair cortisol levels and both intrinsic and ecological traits. Cortisol in blood, feces, and hair are known to be highly correlated in various mammals (e.g. chimpanzees, chipmunks, and mice; Kalliokoski *et al.*, 2019). Therefore, although the concentration values are not directly comparable across different matrices, the effects of covariates can still be compared with our results.

Overall, we found particularly high hair cortisol in most bat species compared to levels reported in hair for other mammals of a similar body size (e.g., root vole [25-50g] = 1.2-6.05; chipmunk [66-110g] = 40.27-260.22 ng/g of hair; Mastromonaco *et al.*, 2014; Książek *et al.*, 2017). Previous studies in bats, examining plasma and feces, have also reported higher cortisol and corticosterone levels relative to similar samples from other mammal species (Klose *et al.*, 2006; Lewanzik *et al.*, 2012; Kelm *et al.*, 2016; Hald, 2019). Some of the exceptional life history traits of bats, such as long lifespan and low fecundity, could explain why bats exhibit higher levels of GCs compared to other mammals (Austad and Fischer, 1991). According to life-history theory, long-lived species with low reproduction rates are expected to prioritize their adult survival (i.e. future offspring) over current reproduction (Stearns, 1992), which could in turn favor higher investment in self maintenance that might be facilitated by high baseline levels of GCs (Ricklefs and Wikelski, 2002).

### Ecological factors among species

Despite the similarity in the HPA hormonal pathways across vertebrates, baseline and stress-induced GC levels are context and species-specific (Romero, 2004; French *et al.*, 2008; Crespi *et al.*, 2013; Kalliokoski *et al.*, 2019). In light of this, it is not surprising that hair cortisol levels in bats showed broad interspecific variation. The differences found could not be explained solely by taxonomic family or phylogenetic relatedness (λ=0), which suggests that other environmental and ecological factors are influencing hair cortisol in Neotropical bats.

Among all the ecological traits evaluated, annual fecundity was the best predictor of hair cortisol. Species with lower fecundity showed higher concentrations of cortisol in hair. We are not aware of any comparable studies systematically examining interspecific variation in cortisol in relation to fecundity in other mammal groups, so we cannot evaluate the generality of our findings in this group. Our results, however, agree with findings from studies in birds, where species with low clutch size and few breeding events also showed higher circulating GCs (Bókony *et al.*, 2009; Ouyang *et al.*, 2011). Life history theory would predict that, for species with lower fecundity, the value of each offspring is higher than in species with relatively high fecundity (Lendvai *et al.*, 2007). Therefore, parents of more valuable broods would be predicted to be more “willing” to invest in offspring survival, which might be facilitated by high baseline GC levels (Bókony *et al.*, 2009). The role of GCs as mediators of the adaptive energy allocation in offspring is complex and requires considering other species-specific aspects such as reproductive strategies, reproductive patterns, and seasonality (Wingfield and Sapolsky, 2003). Data on these factors are currently limited for many Neotropical bat species, making it difficult to properly explore these interactions. Although diet explained additional variation in Neotropical bat hair cortisol, this variable was uninformative; hair cortisol did not vary significantly between our simplified dietary guilds.

Glucocorticoids play a key role in metabolic function, facilitating fuel mobilization (e.g. glucose, fatty acids) under normal and challenging conditions (Kuo *et al.*, 2015). A positive relationship between resting metabolic rate (RMR) and plasma cortisol levels has been reported for various mammalian species (including four species of bats), and this relationship has been suggested as a general pattern for mammals (Haase *et al.*, 2016). Due to the limited data on RMR for our study species, we used basal metabolic rate (BMR) as an indicator of energy expenditure. Different to what we expected, BMR was not an informative factor for cortisol variation in hair among the bats in our study. The positive relationship between cortisol levels and metabolic rate previously found in plasma might be obscured in studies of hair cortisol like ours, due to confounding factors such as moulting cycles and cortisol deposition rate. In addition, obtaining accurate basal metabolic rates in wildlife species (particularly free-ranging animals) is challenging, which raises questions about the quality of BMR data, especially in comparative studies (Genoud *et al.*, 2018). For future studies, a more realistic and informative indicator of energy turnover in free-ranging animals is the Daily Energy Expenditure (Speakman, 1997), which integrates the energy allocated in different activities such as foraging, commuting and thermoregulation (Butler *et al.*, 2004).

The relationship between body condition and GC release has been widely evaluated, because weight loss is one of the early responses to long-term stress in many species (Kitaysky *et al.*, 1999; Angelier *et al.*, 2009; Dickens and Romero, 2013). However, the direction of the effect of body condition on cortisol is context and species-dependent (Crespi *et al.*, 2013). We used body mass as an indicator of body condition because it is a more informative metric than other indexes in bats (McGuire *et al.*, 2018). We found different directions of the effect of body mass on hair cortisol. For three of the studied species (*Molossus nigricans, Pteronotus mesoamericanus* and *Saccopteryx bilineata*), heavier individuals showed higher concentrations of hair cortisol. In contrast, *Pteronotus mexicanus* showed a negative relationship between body mass and cortisol.

One species that stood out for its particularly high levels of cortisol was *Leptonycteris yerbabuenae*. This species is known to be highly mobile and migratory (Horner *et al.*, 1998; Buecher and Sidner, 2013). Migration was not considered in our analyses because the degree to which bats may migrate seasonally is unclear for many of the species in our sample. Migratory behavior, however, could explain such high cortisol concentrations in *L. yerbabuenae*. La Fábrica caves in Colima, one of our field sites, is known to be one of the starting points of the annual migration of *L. yerbabuenae* (Medellin *et al.*, 2018). The role of GCs during migration has been widely studied in birds, fish, and some large mammals, but not in bats (Holberton, 1999; Romero, 2002; Wada, 2008). We hypothesize that premigratory fattening could explain the high hair cortisol levels observed in *L. yerbabuenae*, and we encourage future studies to address this question.

Consistent with other studies, we found differences in hair cortisol levels between sexes, albeit for only four of our 18 studied species: *Desmodus rotundus, Myotis nigricans, Pteronotus mexicanus*, and *Pteronotus mesoamericanus*. For these species, females showed higher cortisol than males, a trend that appears to hold for many mammalian species (Bechshøft *et al.*, 2011; Hau *et al.*, 2016; Rakotoniaina *et al.*, 2017; Dettmer *et al.*, 2018). Higher baseline levels in females can be explained by the differential regulation of gonadal steroid hormones (estrogen and androgens) on HPA axis activity. While estradiol, which is more abundant in females than males, enhances cortisol release, androgens tend to inhibit its production (Handa et al. 1994). Females have also shown differences in HPA axis activity depending on their life history stage, GCs being higher during the late stages of their pregnancy (Reeder *et al.*, 2004). Studies in a fruit-eating bat (*Artibeus jamaicensis*) and little brown myotis (*Myotis lucifugus*) have reported higher levels of plasma GCs in pregnant females (Reeder and Kramer, 2005; Klose *et al.*, 2006). Contrary to their findings, we did not find reproductive state to influence hair cortisol in our female-only model (i.e., for *Pteronotus mesoamericanus* in Belize). However, it may have been difficult to detect an effect given the low number of pregnant females in our sample (n=5, 24%) and the fact that moulting might not occur in conjunction with mating.

Bats have been proposed as good indicators of habitat quality due to their ecological diversity, wide distribution, and potential sensitivity to disturbance (Jones *et al.*, 2009; Stahlschmidt and Brühl, 2012). However, a clear correlation between environmental disturbance and cortisol levels in bats has not been previously reported. Prior studies that examined cortisol in blood did not find differences between bats roosting in agricultural versus urbanized areas (Wada *et al.*, 2010; Allen *et al.*, 2011; Kelm *et al.*, 2016). To assess relationships between disturbance and GCs over longer timescales and without sensitivity to capture stress, we compared hair cortisol in three species found in sites with varying fragmentation and agricultural activities. We found an effect of disturbance in only one of these species, *Pteronotus mexicanus*, which we sampled only in Mexico. Bats roosting in Don Pancho Cave island, a site classified as having intermediate disturbance, showed the highest concentrations of cortisol (Fig.1). We speculate that the high levels found in this population could reflect differences in the cave microhabitat compared to the other caves in our Mexican sample. Don Pancho Cave is a narrow crevice estimated to have a higher colony size (100,000 individuals from 6 species; Téllez *et al.*, 2018) than the other sampled caves El Salitre (~10,000 individuals from 10 species; Torres-Flores *et al.*, 2012) and La Fábrica (>5000 individuals from four species). The higher density of bats living in the Don Pancho cave may mean that there are increased agonistic social interactions in this population (Creel *et al.*, 2013), that the risk of parasite exposure is increased (Postawa and Szubert-Kruszyńska, 2014), and parasite transmission rates are higher (Langwig *et al.*, 2012). Based on studies in other mammals, all of these factors might explain the high cortisol levels found in the Don Pancho Cave population of *P. mexicanus*; however, the influence of these factor on hair cortisol in bats is still unknown.

Physiological responses to chronic stress in wildlife are difficult to unravel and predict unless multiple responses at different levels of biological organization are evaluated simultaneously (Dickens and Romero, 2013). Hair cortisol offers great potential as a tool to monitor health in wild populations, particularly those already identified at risk (Kalliokoski *et al.*, 2019). For instance, chronically elevated cortisol levels have been linked to greater susceptibility to infection and disease severity (Davy *et al.*, 2017).Periodic surveys of hair cortisol could therefore help identify periods when bats might be more vulnerable to infection (e.g. white nose syndrome). Further, such surveys might also inform when individuals are more likely to shed zoonotic pathogens (e.g. henipaviruses and filoviruses; Plowright *et al.*, 2008; Davy *et al.*, 2017; McMichael *et al.*, 2017; Kessler *et al.*, 2018).

## Conclusions

The current study reports cortisol levels in hair of 18 Neotropical bat species from two countries and serves as a reference for future research using this method in wild bat populations. We found that fecundity and potentially diet are important ecological traits explaining interspecific variation in bat hair cortisol. Within species, female bats exhibited higher cortisol than males, and the effect of body mass varied among species. Other factors that may be important at the individual level, such as parasite load and colony size, should be considered in future studies to have a more complete understanding of sources of variation on baseline GC levels within species. Importantly, studies looking at hair growth rate and moulting cycles in Neotropical bat species are imperative to give an accurate interpretation of hair cortisol as a biomarker of stress response. Applied properly, hair cortisol quantification is a powerful non-invasive technique with multiple potential applications in bat ecology, physiology, and conservation. Our findings and ongoing work will help to validate and apply hair cortisol as a monitoring tool in wild bat populations.

## Acknowledgments

We thank Christine Gilman, Patricia Medd, and Paula Mackie for their assistance with the cortisol assays. We also thank Brock Fenton, Sara Ketelsen, Neil Duncan, the numerous members of the Lamanai bat research team, and the staff of the Lamanai Field Research Center for their assistance with bat capture, field logistics, and permits in Belize.

## Funding

This work was supported by the Natural Sciences and Engineering Research Council of Canada Discovery Grant [#386466] to KCW. GM was supported by the Toronto Zoo Foundation (GM), DJB was funded by the ARCS Foundation and the American Museum of Natural History Theodore Roosevelt Memorial Fund, and NBS was supported by the Taxonomic Mammalogy Fund of the American Museum of Natural History.

## Supplementary material

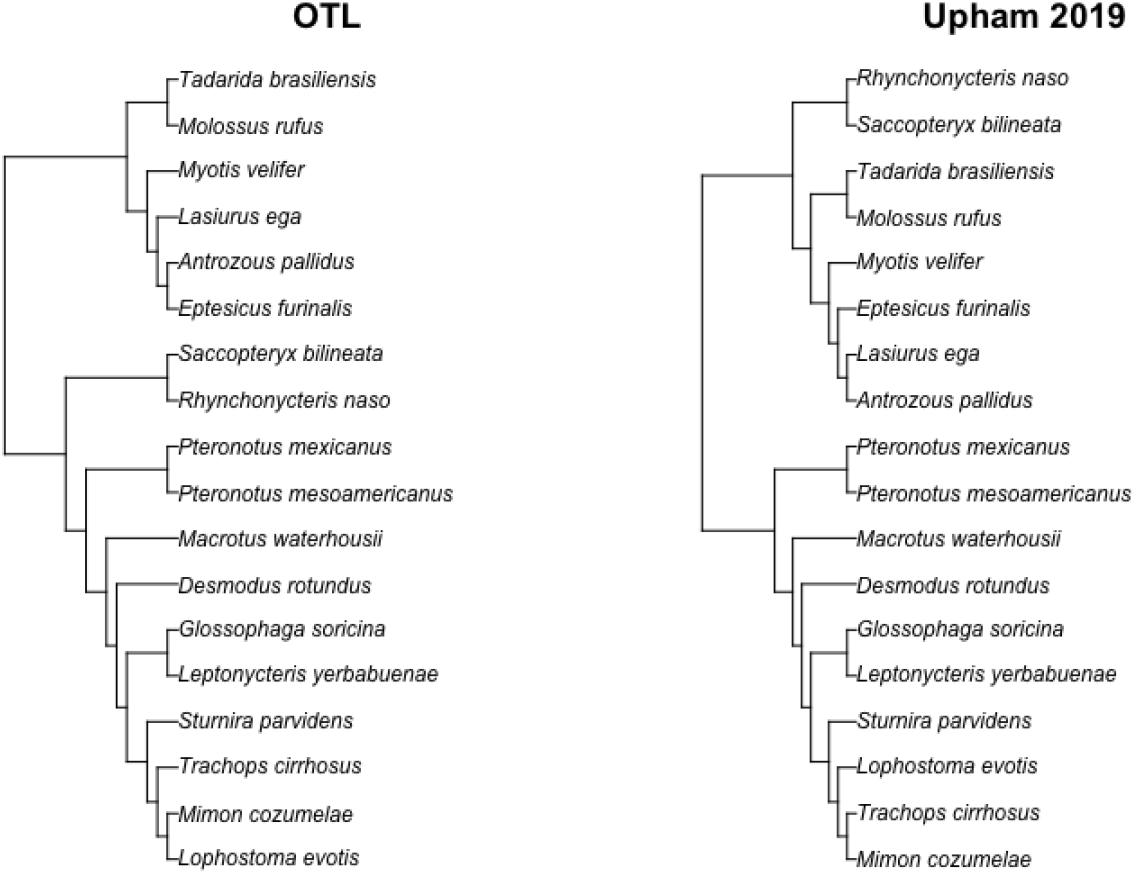

## Notes

### Competing Interest Statement

The authors have declared no competing interest.

### Summary of Updates

Corrected error in number formatting and units for the values presented in the third line of the results sections.

